# A Transcriptome Fingerprinting Assay for Clinical Immune Monitoring

**DOI:** 10.1101/587295

**Authors:** Matthew C Altman, Nicole Baldwin, Elizabeth Whalen, Taha Al-Shaikhly, Scott Presnell, Prasong Khaenam, Vivian H Gersuk, Laurent Chiche, Noemie Jourde-Chiche, J Theodore Phillips, Goran Klintmalm, Anne O’Garra, Matthew Berry, Chloe Bloom, Robert J Wilkinson, Christine M Graham, Marc Lipman, Ganjana Lertmemongkolchai, Farrah Kheradmand, Asuncion Mejias, Octavio Ramilo, Karolina Palucka, Virginia Pascual, Jacques Banchereau, Damien Chaussabel

## Abstract

**Background:** While our understanding of the role that the immune system plays in health and disease is growing at a rapid pace, available clinical tools to capture this complexity are lagging. We previously described the construction of a third-generation modular transcriptional repertoire derived from genome-wide transcriptional profiling of blood of 985 subjects across 16 diverse immunologic conditions, which comprises 382 distinct modules.

**Results:** Here we describe the use of this modular repertoire framework for the development of a targeted transcriptome fingerprinting assay (TFA). The first step consisted in down-selection of the number of modules to 32, on the basis of similarities in changes in transcript abundance and functional interpretation. Next down-selection took place at the level of each of the 32 modules, with each one of them being represented by four transcripts in the final 128 gene panel. The assay was implemented on both the Fluidigm high throughput microfluidics PCR platform and the Nanostring platform, with the list of assays target probes being provided for both. Finally, we provide evidence of the versatility of this assay to assess numerous immune functions *in vivo* by demonstrating applications in the context of disease activity assessment in systemic lupus erythematosus and longitudinal immune monitoring during pregnancy.

**Conclusions:** This work demonstrates the utility of data-driven network analysis applied to large-scale transcriptional profiling to identify key markers of immune responses, which can be downscaled to a rapid, inexpensive, and highly versatile assay of global immune function applicable to diverse investigations of immunopathogenesis and biomarker discovery.

## BACKGROUND

Traditionally the immune system has been viewed as playing a beneficial role in control of infection and a detrimental role in autoimmunity and allergic processes. More recently it has been appreciated to also have critical functions in a far wider range of common diseases including obesity, atherosclerosis, dementia, and numerous cancers among others[1-4]. The immune system acts as a highly interconnected network of cellular and humoral interactions and it is through either appropriate function or malfunction of network components that the immune system underpins these diverse human diseases [5-9].

A better understanding of individual immune processes has facilitated numerous interventions that can alter immune responses and ameliorate outcomes. However cost-effective and standardized tools available to practitioners and clinical trialists that capture the complexity of immune responses and that could be used for monitoring the immune status of patients are lacking. Global monitoring of the immune system, even in clinical trials of immunotherapies, is often missing due to the complexity of implementing systems approaches.

Immune responses are highly complex, dissecting out the reproducible global patterns of the immune system is critical to developing improved methods of immune monitoring [10]. In this regard, systems-scale analyses of global architecture in both normal and pathologic immune function is necessary. Different technologies under development and used in research have numerous strengths but also distinct challenges, including cost of reagents or instruments, complexity of the assay workflow, and complexity of data analysis and interpretation.

We previously described the construction of a third-generation modular repertoire, compromised of 382 modules, and is representative of 16 immune states. We also described transcriptome fingerprinting as a novel and useful visualization scheme [11]. Here we take our work a step further, and describe the development of a cost-effective and practical, targeted transcriptome fingerprinting assay (TFA) (**Figure 1**), while preserving its capability of monitoring the same modular repertoire of immune responses. This assay is by design meant as a generic assay suitable for immune profiling across multiple states of health and disease. We demonstrate the utility of creating such an assay through a purely data-driven network analysis approach to identify core functional immune pathways. Our results show the complex molecular interactions in immune pathogenesis but also reveal a redundancy of core immune circuits.

**Figure 1:**
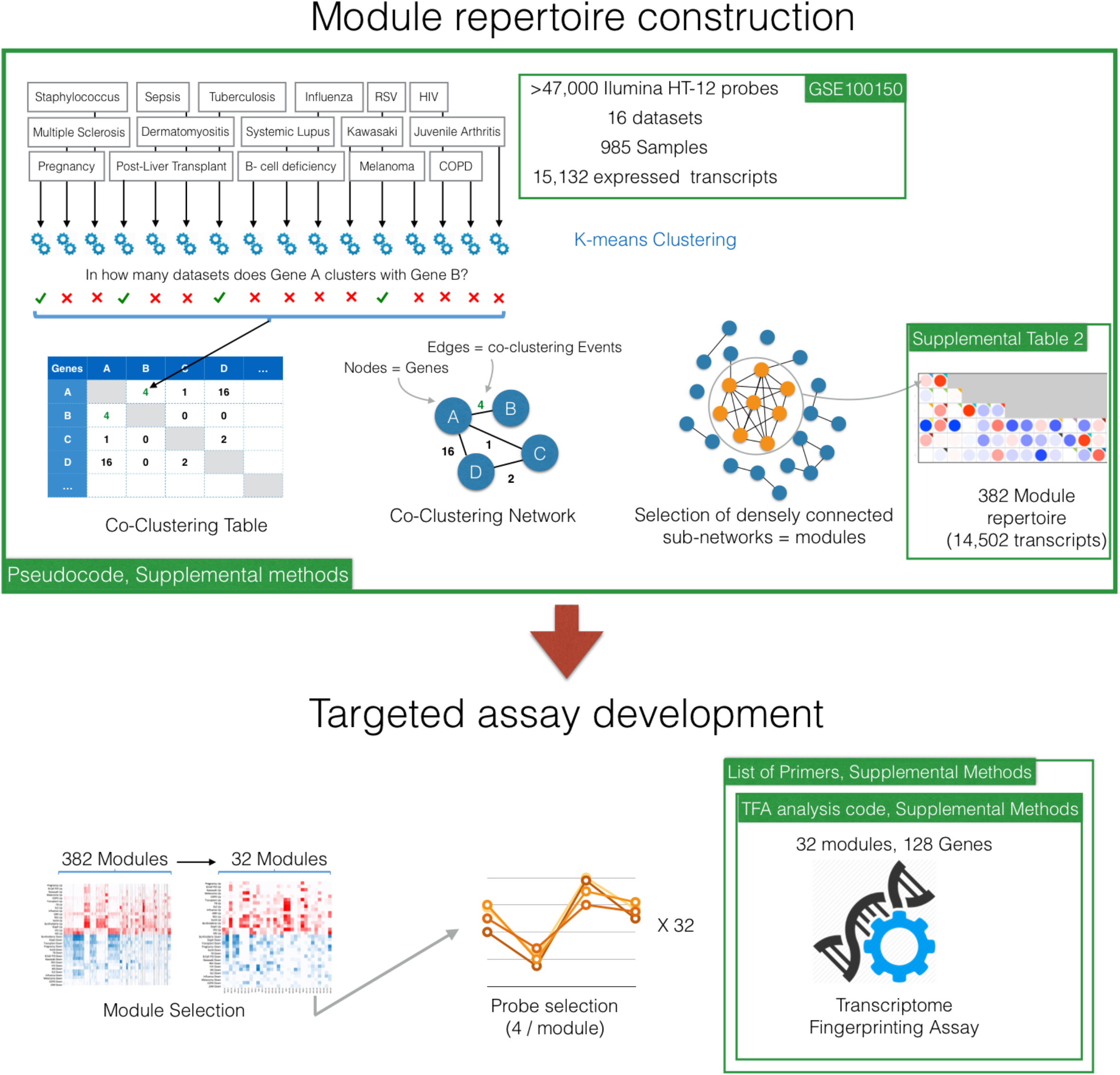
Overview of the module repertoire construction and targeted assay development approach. All details are provided in the main text and supplemental methods sections. Briefly, blood transcriptional module repertoire construction takes a collection of transcriptome datasets as input. In this case 16 datasets constituted by 985 individual transcriptome profiles spanning a wide range of immunological and physiological states. Clustering behavior of gene pairs is recorded for each independent datasets and the information complied in a co-clustering table. Subsequently the co-clustering table serves as input for the generation of a co-clustering graph, where nodes are the genes and edges represent co-clustering events. Next the largest, most densely connected subnetworks among a large network constituted of 15,132 nodes are identified mathematically and assigned a module ID. The genes constituting this module are removed from the selection pool and the process is repeated. The resulting framework of 382 modules served as a basis for the development of targeted assay. This involves two major steps. First, the selection of representative modules among the 382 modules constituting the framework. Second, the selection of representative probes among those modules. The process can be adjusted according to practical constraints, such as assay throughput and cost. In our case the selection of 32 modules out of the original set of 382, and of 4 representative genes from each of the 32 modules yielded a 128-gene fingerprinting assay.

This assay highly simplifies the analysis and interpretation of gene transcription data in immunologic diseases. Several applications are envisioned, including assessment of disease activity, health monitoring, pre-symptomatic detection of disease, and biomarker discovery. In this manuscript we show application of the assay to assess disease activity in systemic lupus erythematosus (SLE) and for longitudinal immune monitoring during pregnancy. This assay is released as an “open resource”, including a complete list of reagents, detailed procedures, and source code for data analysis.

## RESULTS

In our third-generation modular repertoire, the 382 modules showed an inherent variability. We sought to show that the variability apparent in the modules can be reduced by using a subset of the module genes, demonstrating that there are core genes within each module that best reflect perturbations of that pathway. It also produces sets of genes that support rapid and cost effective transcriptional profiling. In order to find a representative subset of modules that best reflects the variability seen across the source data, we partitioned the modules with Hartigan’s K-means algorithm using the jump statistic [12] to determine an appropriate number of clusters and to reduce granularity, resulting in 38 subgroups. The module closest to the mean vector in each subgroup was selected to represent that subgroup, and if a subgroup did not contain at least one module in one of the 16 diseases showing at least 25% of genes up-or down-regulated, it was excluded from selection. This left 32 modules representative of the 382 original modules (**Figure 2**). From each of the 32 modules we then selected 4 representative probes by ranking all probes according to the distance of each probe from the module’s mean probe vector. The highest ranking probes that had gene symbols unique to a module were selected.

**Figure 2:**
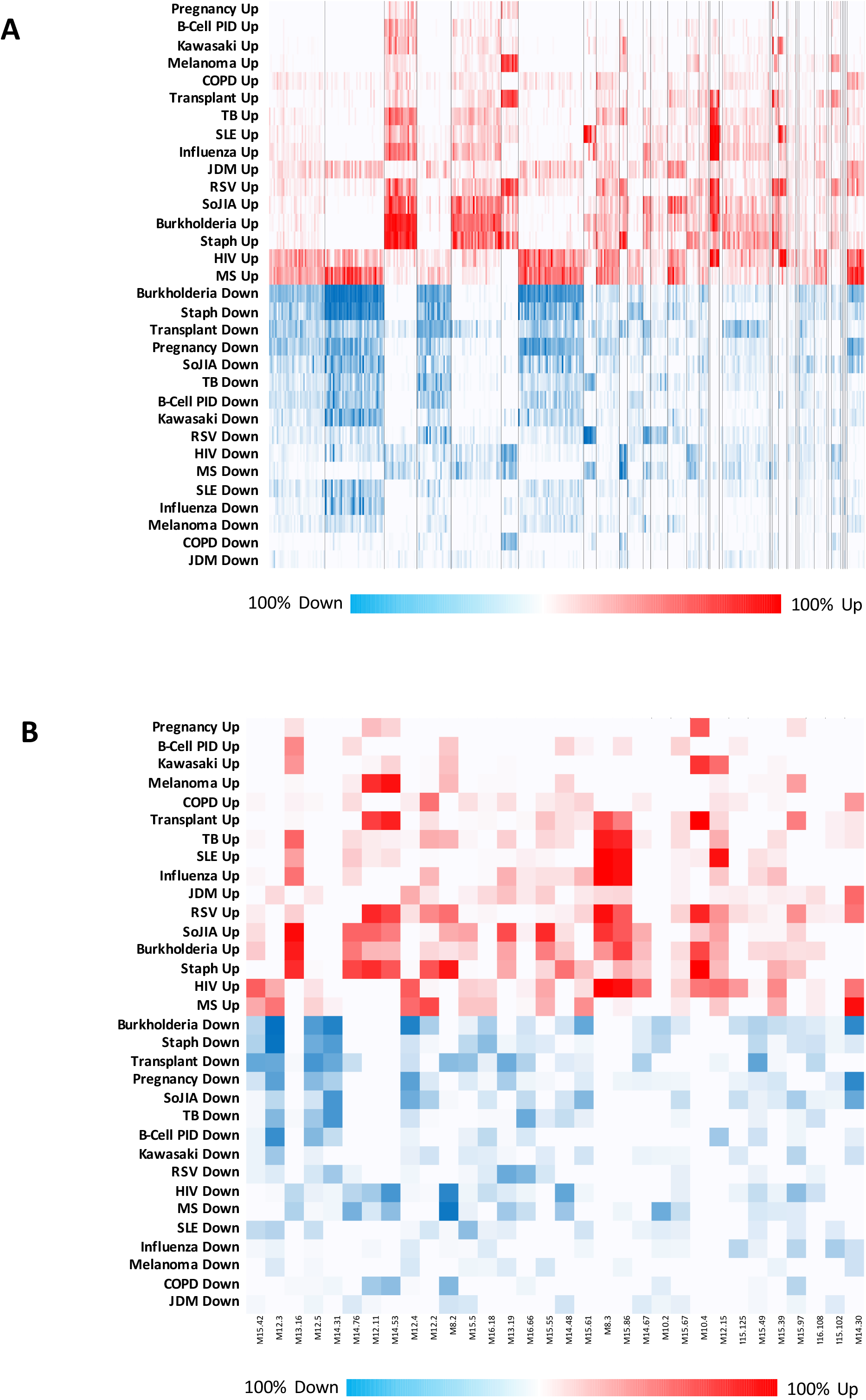
Patterns of blood transcript abundance observed across 16 disease or physiological states. **(A)** Depicted is the expression pattern of the gene members in each of the 382 modules (columns) across all 16 disease states (rows). Each pixel represents the percent of probes within that module that show a significant difference in expression between the disease group and the control group within that microarray dataset. The color scale ranges from 100% up-regulated (full red) to 100% down-regulated (full blue). The modules are clustered into 38 distinct subgroups separated by black vertical lines according to similarity of expression pattern across the 16 disease states. **(B)** The expression pattern of the gene members in each of the 32 representative modules of the 382 original modules/38 clusters depicted in **(A)** representing the downscaled 32 modular repertoire.

This unbiased process selected out modules most representative of each cluster and genes most representative of each module, resulting in a subset of 32 modules and 4 genes per module for a total of 128 genes representative of the diversity among the 382 modules and 14,502 total transcripts profiled. These 32 modules/128 genes were chosen to represent the “transcriptome fingerprint” used for assessment and monitoring of immunologic disease states (Table 1). This scaling down process allows the implementation of targeted assays, which, given the markedly reduced volumes of data generated per experiment, are cost-effective and allow higher sample throughput, faster turnaround and streamlined analysis and interpretation, while maintaining profiling of a broad repertoire of immune gene signatures.

**Table 1:**
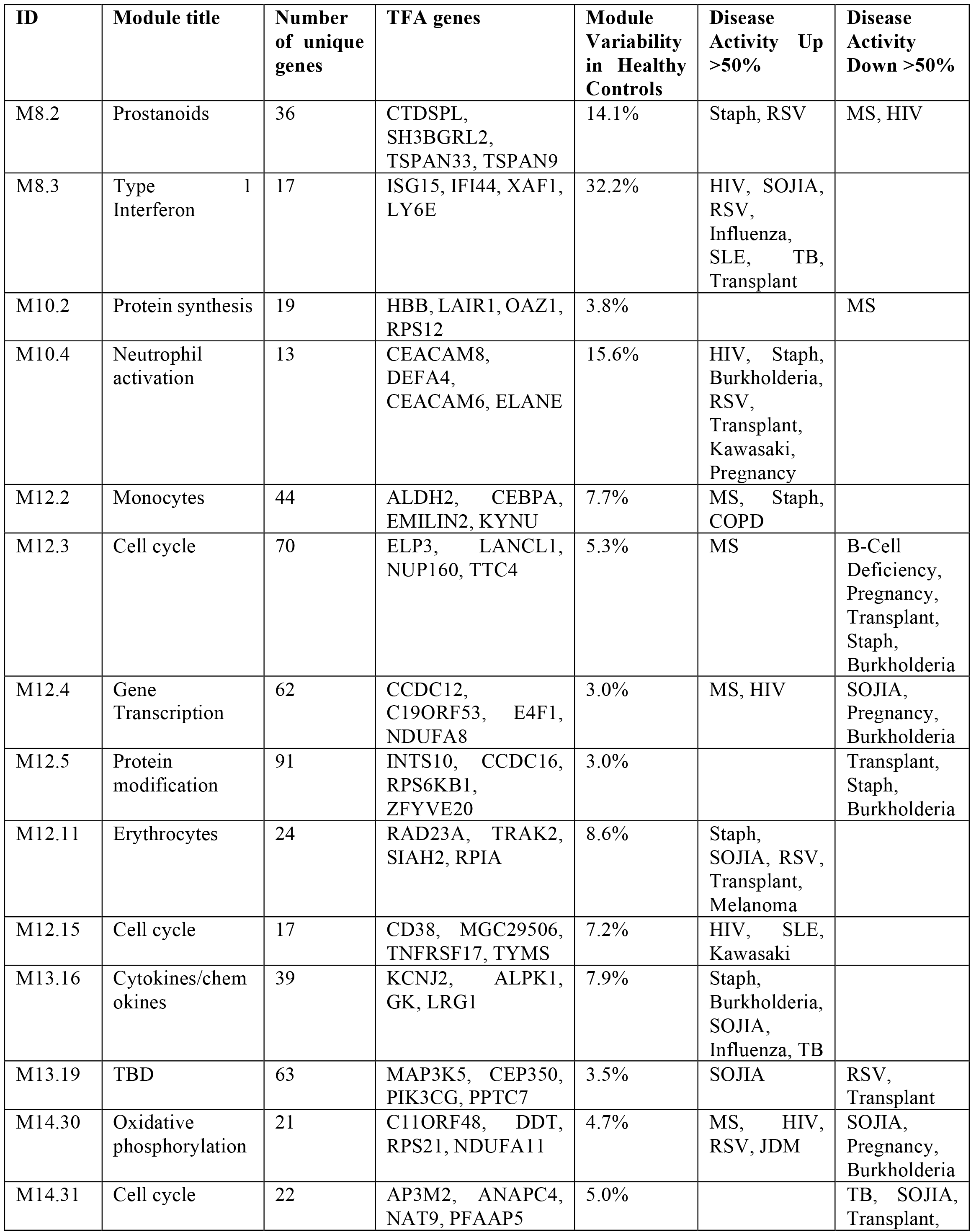
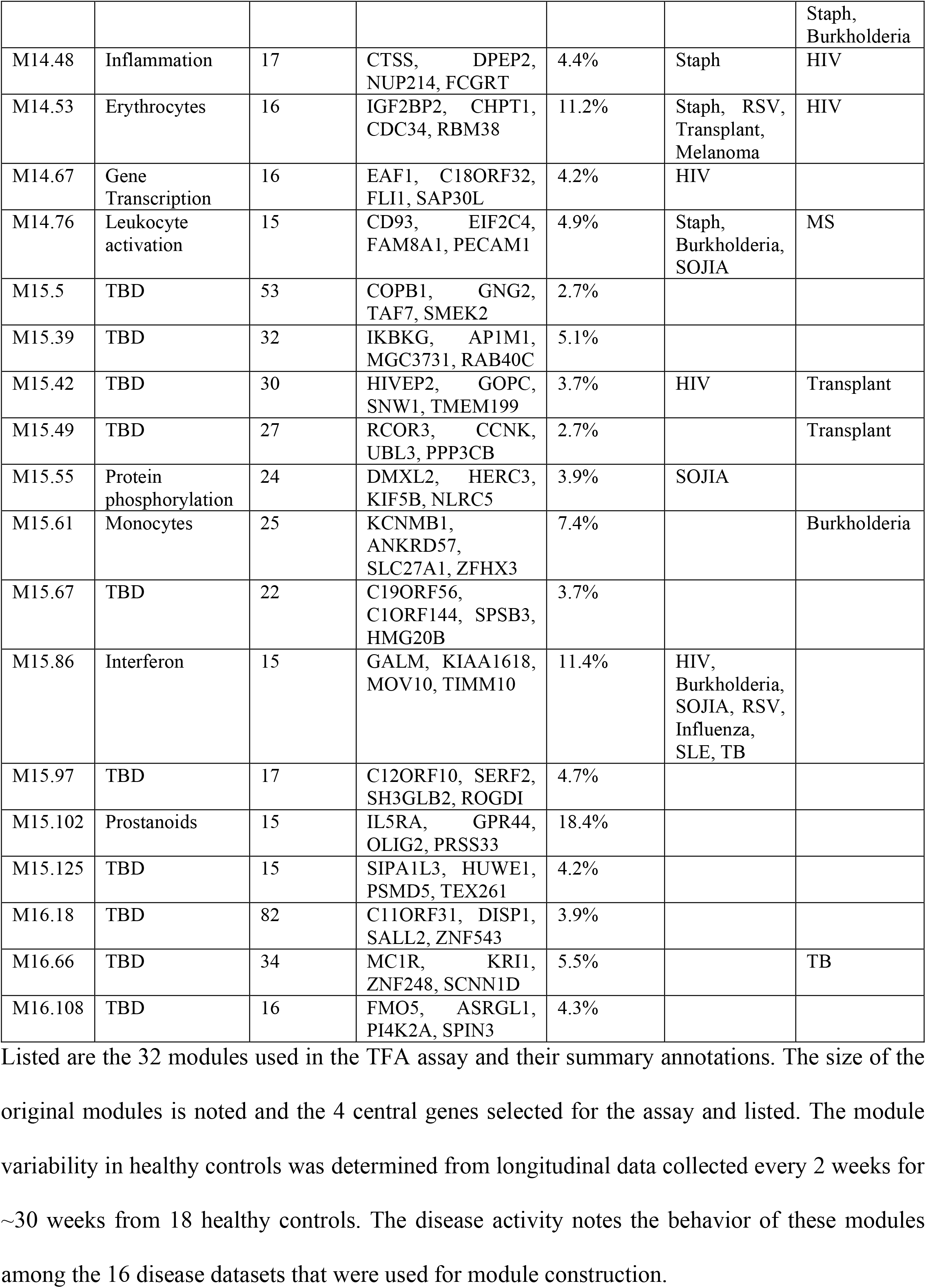
TFA modules.

### Identification of biomarkers for disease activity in SLE

We first sought to show that a transcriptome fingerprinting assay (TFA) panel when measured by high-throughput qPCR, shows high correlation with gene expression data derived from microarray. For this we used a cohort of adult patients with SLE, which was independent from the cohort used for module construction [13]. Whole blood samples from 24 SLE patients with varying stages of disease activity as well as 15 healthy age and gender matched controls were used for analysis. Samples were selected that had been drawn prior to initiation of new immunosuppressive treatment to reflect disease activity before therapy. Whole blood derived RNA samples taken at an initial visit were used to generate both microarray and qPCR data. High correlation values were observed between microarray and qPCR data for all genes and modules that had FC values greater than 1.5 for the ratio between SLE and healthy control expression levels (**Figure 3a**).

**Figure 3:**
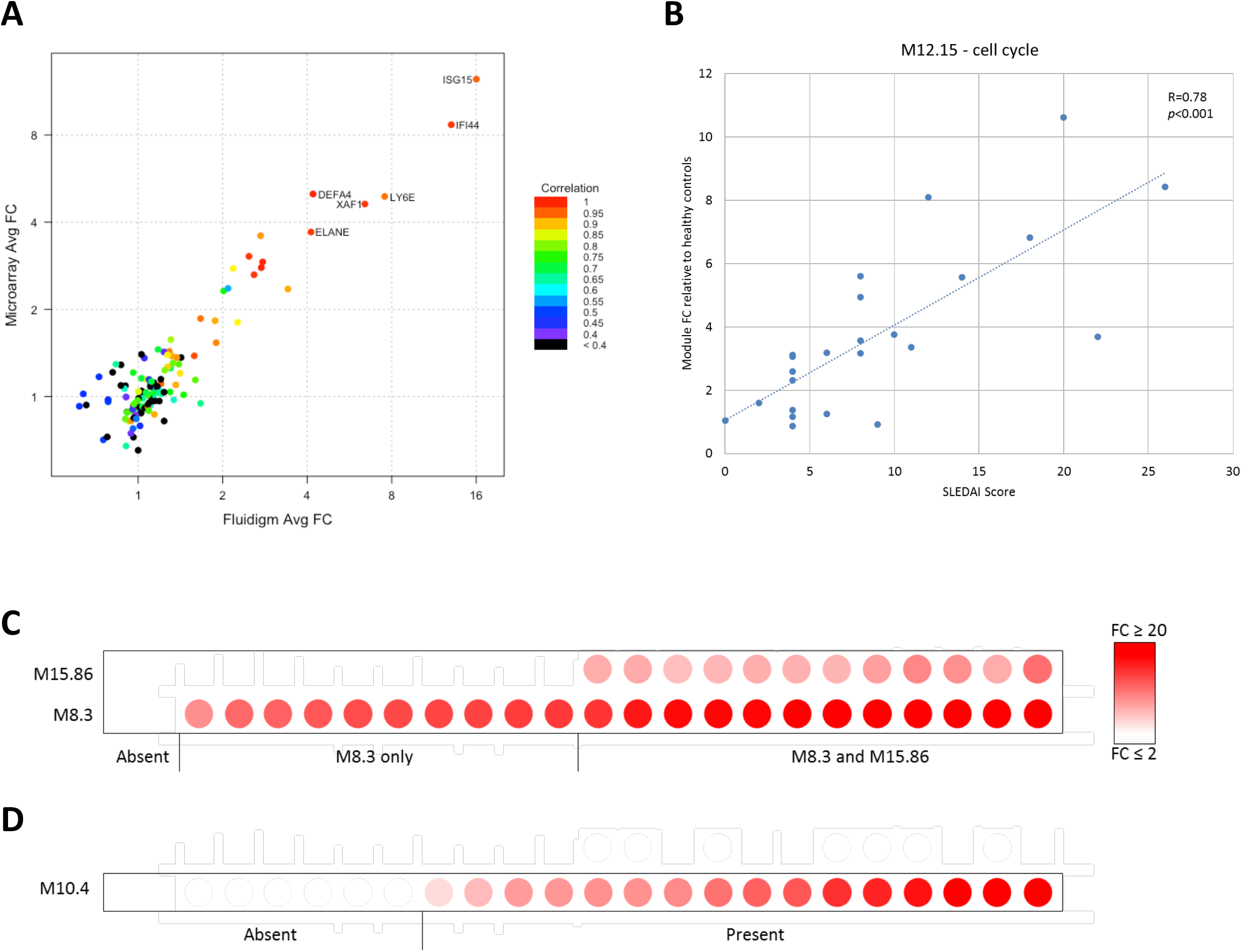
TFA analysis of SLE demonstrates activation of cell cycle, interferon, and neutrophil pathways. **(A)** Each point represents the fold change of a single gene comparing the average of SLE patients to the average of Controls as measured by qPCR (x-axis) and microarray (y-axis). Average fold change values were similar and proportional across technologies. The color of each point represents the Pearson’s correlation of fold change values among SLE patients for a given gene as measured by qPCR and by microarray. Those genes with high average fold change values also showed very high levels of correlation across platforms. **(B)** Fold change value of module M12.15 (average of 4 genes) plotted versus the SLEDAI score for each SLE patient. Correlations and *p* values were calculated using Spearman’s rank correlation coefficient. **(C)** Interferon module fold change values are shown for each SLE patient (n=24) compared to the average of the healthy controls. Each point represents the module fold change of a single SLE patient. Patients are ordered according to increasing FC of M8.3. The greater the intensity of each point, the greater the fold change. White FC≤2. Full red ≥20. **(D)** Neutrophil module fold change values are shown for each SLE patient (n=24) compared to the average of the healthy controls. Each point represents the module fold change of a single SLE patient. Patients are ordered according to increasing FC of M10.4. The greater the intensity of each point, the greater the fold change. White FC≤2. Full red ≥20.

Next, differences in immunologic pathways potentially useful for stratifying the SLE patients were evaluated. Four modules that are biologically relevant to SLE showed distinct up-regulation in SLE relative to healthy controls. These were M12.15, annotated as a cell cycle module, M8.3 (type 1 interferon), M15.86 (interferon), and M10.4 (neutrophil activation). Module M12.15 (cell cycle) showed a high degree of correlation with clinical disease activity as measured by the SLE Disease Activity Index (SLEDAI) score (R=0.793, P=6.03E-6) [14]. This module showed a higher level of correlation than traditional markers of disease activity – anti-double-stranded DNA titers (R=0.636, P=0.001) and C-reactive protein (CRP) (R=0.327, not significant) – which had been measured at the same time (**Figure 3b**) [15]. The four representative genes within module M12.15 are linked to SLE. TYMS is an enzyme critical for folate metabolism and polymorphisms in this and related genes have previously been associated with SLE [16]. CD38 has been shown to be expressed in higher levels on circulating lymphocytes in active SLE [17]. TNFRSF17 (also called BCMA) is a B-cell maturation antigen that has been shown important for B-cell development and SLE pathogenesis in mouse models [18]. MZB1 (previously MGC29506) has been shown to be involved in immunoglobulin heavy chain biosynthesis [19]. Our finding demonstrates that four genes representative of a signature important to B cell development show very high correlation with SLE disease activity and that TFA could present potential clinical utility for assessment of disease activity.

Abundance of transcripts representative of two interferon modules, M8.3 and M15.86, was also increased in the SLE samples relative to control. Several groups have previously reported an increase in type I interferon (IFN) regulated genes in a subset of SLE patients (reviewed in [20] and [21]). More recently, whole-genome transcriptional profiling has demonstrated a higher degree of complexity in interferon activity with an apparent gradient of interferon responses in SLE [13, 22]. Here our findings support this observation; since patients with SLE included in our study could be similarly stratified based on either absence of interferon signature, activation of a single interferon module M8.3, or activation of both interferon modules M8.3 and M15.86, in an apparent sequential pattern (**Figure 3c**). The genes representative of M8.3, ISG15, IFI44, LY6E, and XAF1, are all well characterized as IFN-alpha inducible genes. The genes representative of M15.86, MOV10, TIMM10, KIAA1618, and GALM are less well characterized and may represent either a related or distinct interferon response seen in a subset of SLE patients in parallel with a relatively saturated IFN-alpha response. TFA provides a straightforward framework to further investigate this differential interferon response in SLE.

Module M10.4 is composed of neutrophil specific genes, but did not show significant correlation with participants’ absolute neutrophil counts collected at the same time (R=0.33, P=0.144), suggesting this module reflects alterations in neutrophil function and activity more than quantity. The four representative genes that were selected out of the fifteen genes constituting M10.4 encode well characterized proteins important to neutrophil function; CEACAM6 and CEACAM8 are cell-adhesion proteins on neutrophils essential for neutrophil adhesion and migration [23]; DEFA4 is a defensin peptide, and ELANE or neutrophil elastase is a serine protease, both are contained in neutrophil azurophil granules. Many, but not all patients show significant increase in abundance of transcripts representative of M10.4, suggesting abnormal neutrophil function in only a specific subset of untreated SLE (**Figure 3d**). There is growing evidence for alterations in neutrophil chemotaxis, phagocytosis, superoxide production, and apoptosis in subgroups of SLE patients and investigation of neutrophil activity is an active area of research in SLE pathogenesis[24-26]. Our assay agrees with findings that neutrophil dysregulation may be observed in only a specific subset of SLE and suggests that this gene set could serve as a marker to identify SLE patients with neutrophil dysregulation.

### Longitudinal monitoring of immune status in pregnancy

To determine the baseline expression of TFA modules in healthy adults and to assess their utility for monitoring of immunologic changes, the assay was run on samples collected longitudinally from 18 healthy non-pregnant volunteers and 12 healthy pregnant women. Samples were collected at 2-week intervals for up to 28 weeks. In the pregnant women, sample collection started at ∼10 weeks into the pregnancy. All of these women had uncomplicated term deliveries of healthy infants. We monitored changes in gene expression of these 32 modules over time to assess for consistent changes attributable to pregnancy which is likely to mediate progressive immunological and physiological changes over time.

Investigation of the healthy controls demonstrated no significant differences in expression levels of these 128 genes according to gender, age, or time point (**Supplemental Figure 1**). Therefore, these 18 individuals were used to define reference ranges for expression of these 128 genes. To support comparisons amongst modules, all expression values were scaled to a mean of 1, and confidence intervals defined by the standard errors of the healthy controls. Most modules (26/32) had narrow confidence intervals of +/- 10% or less, demonstrating these genes show relatively modest fluctuation in healthy individuals (Table 2). Only one module, M8.3 (type 1 interferon), showed particularly high variability (+/- 32%), which we believe could be related to viral infections in some of the controls throughout the course of monitoring or else may reflect higher normal temporal variability of expression of these genes.

Samples from the pregnant women were tested for consistent changes during the course of monitoring as well as group differences compared to healthy controls. After multiple testing correction, 7 modules, M10.4, M8.2, M12.11, M14.53, M13.16, M12.2, and M15.55, showed a significant linear change over time. Another 4 modules, M14.76, M15.102, M12.4, and M12.3, were significantly different from controls over the period of time monitored suggesting a change in expression occurred in the first 10 weeks of pregnancy prior to the start of sample collection (**Figure 4** and **Supplemental Figure 3**).

**Figure 4:**
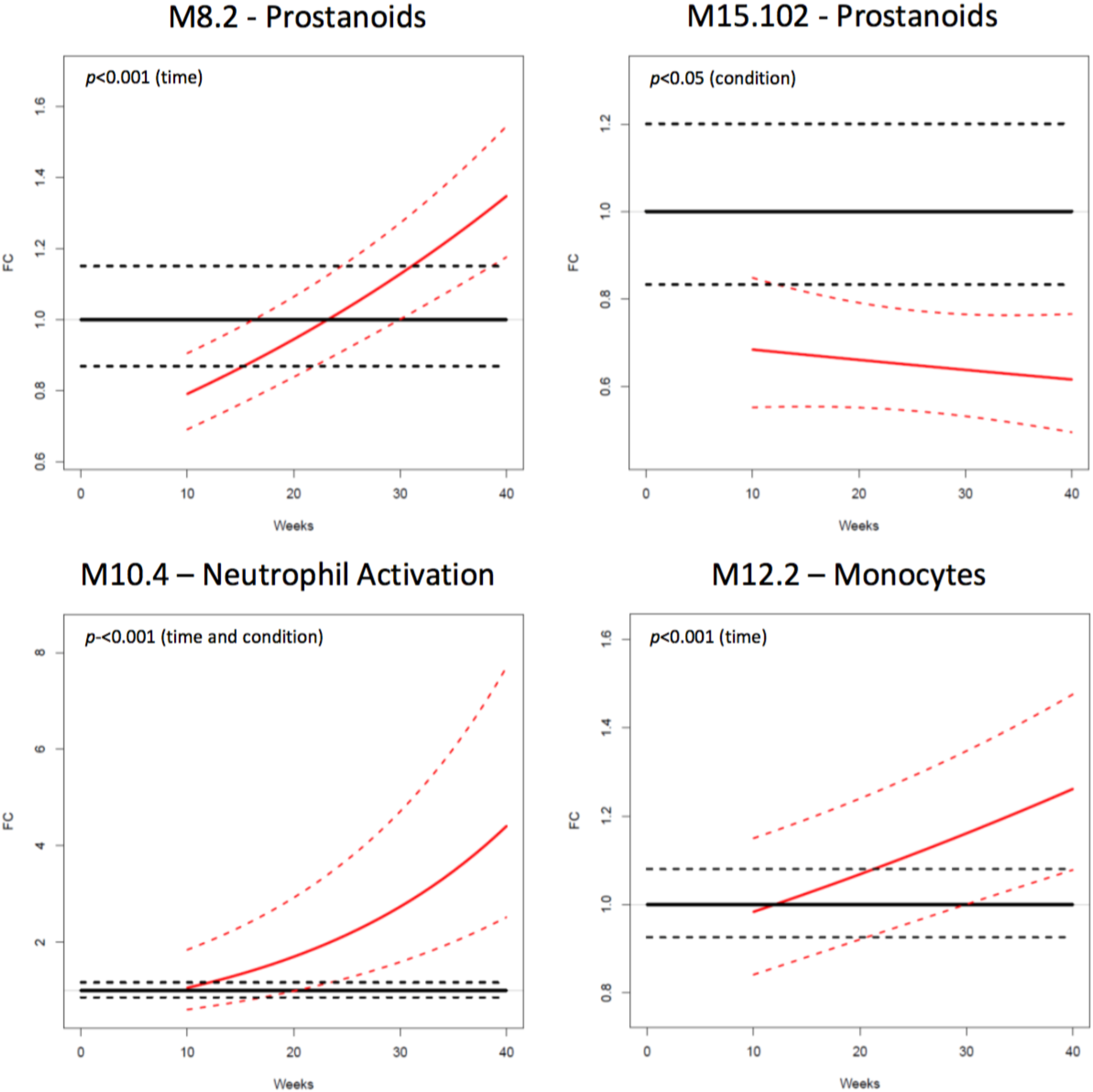
Changes in blood transcript abundance measured during the course of uncomplicated pregnancies. Average FC expression levels referenced to healthy controls from the blood of 12 healthy pregnant women and 18 healthy non-pregnant controls. Shown are 4 of the 11 significant modules (see **Supplemental Figure 3** for the other 7). A linear mixed effects model was fit to the longitudinal data from pregnant women and healthy controls to determine if there was a trend over time for the pregnant women (time *p*-value) and if there was a significant difference between pregnant women and healthy controls over time (condition *p*-value).

Several of these modules represent immune functions that are known to change during healthy pregnancy. M8.2 and M15.102 are both relevant to prostanoid metabolism. Prostanoids are critical for cervical and uterine development in preparation for delivery [27]. M8.2 shows a linear increase in expression throughout pregnancy with fairly narrow confidence intervals (+/- 13%) and based on the genes in M8.2 likely relates to increasing prostanoid production. M15.102 is decreased relative to controls throughout monitoring. M15.102 includes both GPR44 (also called PTGDR2) a prostaglandin D2 receptor and IL5RA a subunit of the IL5 receptor, both of which play critical roles in T helper cell mediated immune responses, as well as OLIG2 and PRSS33, which are less well characterized. The decrease in this module may reflect a change in lymphocyte function in relation to changing levels of prostanoids.

M10.4 is a neutrophil activation module. It increases dramatically during pregnancy to more than 4-fold on average. This is likely in part due to the known increase in neutrophil numbers in the peripheral blood during the 2nd and 3rd trimesters, but likely also reflects changes in neutrophil function as previously observed [28] since the genes composing this module are specific to neutrophil activation as discussed earlier. Module M12.2 (composed of genes ALDH2, CEBPA, EMILIN2, and KYNU) is a monocyte module. Monocytes generally do not change in number during pregnancy but show increased activation in the circulation [29]. The 7 other modules that show different expression patterns compared to healthy controls represent several other biological processes including novel findings that compel further investigation (**Supplemental Figure 2**). Taken together our findings show that the TFA assay results are stable over time in non-pregnant healthy adults and can detect progressive immunologic changes during the course of a healthy pregnancy. This provides a baseline for further investigation of immunologic changes that can occur during both healthy and complicated pregnancies.

## DISCUSSION

We present here, the design and implementation of modular transcriptional repertoire-based targeted assays. It is based on our prior work in constructing a third-generation modular repertoire in clinical immunology, whereby variation in abundance of blood RNA was captured through the construction of co-clustered transcriptional modules. The modular repertoire, was representative of 16 immune states (16 instead of 7 and 8 in earlier generations) [30, 31], and used as an input transcriptome profiles of nearly 1000 subjects. That approach identified 382 modules showing co-clustering across a wide range of immune conditions, while others appeared to be more condition-specific. In this work, we show that modules can be reduced to representative genes in a purely data-driven fashion that does not depend on a priori knowledge about the genes or clinical states.

Using such approach, we found that useful representative genes of a functional pathway may not be canonical genes, and that gene selection through a data-driven network analysis approach is powerful for novel discovery and assay development. This method, which we have called a “transcriptome fingerprinting assay” or TFA, enabled down-scaling from complicated genome-wide expression profiling to rapid and cost-effective qPCR and molecular barcoding platforms. Proof of principle was provided for disease pathogenesis, biomarker discovery, and longitudinal monitoring applications. TFA was employed to investigate immune perturbation in SLE and pregnancy. We were first able to establish the high degree of correlation between TFA and microarray data, and to demonstrate stability of the TFA gene signature in healthy adults over time. More importantly we were also able to demonstrate the ability of this assay to detect both known and novel biological changes. In the case of SLE, confirming for instance the differential expression of interferon genes, and adding evidence regarding neutrophil dysregulation in a subset of patients. We also found that a cell cycle module shows a very high degree of correlation with SLE disease activity, which warrants further investigation as a potential disease biomarker. Clinical utility may be found through combination of these modules to provide rapid and effective means to stratify and monitor SLE patients.

In the setting of pregnancy, numerous modules involved in prostanoid metabolism, neutrophil activation, and monocyte activation were found to change in a coherent fashion throughout the course of the second and third trimester. Some of those modules constitute a means to quantify immune changes that are known to take place during pregnancy while others appear to track changes that have not previously been recognized and that will need to be further characterized. Indeed, currently there are no biomarkers for two of the most common adverse pregnancy outcomes: preterm labor/delivery and preeclampsia/eclampsia [32, 33]. Risk factors for preterm labor and delivery are believed to act through multiple immune pathways including altering eicosanoid metabolism and increasing prostaglandin production [34, 35] and through changes in neutrophil cytokine production [36]. Similarly changes in eicosanoid metabolism play a major role in preeclampsia and eclampsia [37], and both neutrophil and monocyte activation are thought to be important in the pathophysiology of this condition [28, 38, 39]. We hypothesize that longitudinal measurement using TFA modules, in particular those we have demonstrated to have coherent change throughout pregnancy, could be used for a better understanding and eventually early detection of maternal and perinatal complications such as preterm birth and pre-eclampsia. Testing of this hypothesis is currently underway.

## CONCLUSION

To conclude, this work demonstrates the utility of purely data-driven network analysis applied to large-scale transcriptional profiling datasets to identify key markers of immune responses. From this approach we have developed a transcriptome fingerprint of the immune system based on a non-systems scale assay, which is applicable to investigation of immunopathogenesis, longitudinal monitoring, and biomarker discovery. Sample acquisition for this assay is straightforward as blood can be collected by venipuncture or finger stick [40] and requires no onsite processing. The TFA assay is cost-effective, generates a manageable volume of data, and does not require sophisticated bioinformatics infrastructure and pipelines for analysis. Notably the successful use of both a PCR based assay and a well-established molecular barcoding technology (NanoString) confers additional advantages, since both technologies are known for sensitivity, robustness, and ease of use. Furthermore PCR is widely used in clinical diagnostic and research settings, which would allow our assay to be easily adopted. We are publishing this assay as an “open resource”, including a complete list of reagents and source code for data analysis. This should facilitate third party implementation of the assay and hopefully encourage re-sharing of iterative improvements of its design and of the downstream analytic pipeline. Indeed, taken together, the development of streamlined “Omics-based” assays should contribute to a wider adoption of systems approaches, or in this case “systems-based” approaches.

## METHODS

### Modules/Genes Selection - Downscaling to transcriptome fingerprinting

The 382 modules comprising our third generation modular repertoire were grouped using Hartigan’s K-means algorithm and using the jump statistic to determine an appropriate number of clusters [12], resulting in 38 subgroups. The module closest to the mean vector in each subgroup was selected to represent that subgroup. If a subgroup did not contain at least one module in one of the 16 diseases showing at least 25% of genes up-or down-regulated, it was excluded from selection. This left 32 modules representative of the 382 original modules. From each of those 32 modules, 4 representative probes were selected by ranking all probes according to the distance of each probe from the module’s mean probe vector and the number of presence calls per sample group (detection P< 0.01). The highest-ranking probes that had gene symbols unique to a module were selected.

### Microarray data generation (SLE cohort)

Globin mRNA was depleted using the GLOBINclear^™^ (Thermo Fisher Scientific). Globin-reduced RNA was amplified and labeled using the Illumina TotalPrep-96 RNA Amplification Kit (Thermo Fisher Scientific). Biotin-labeled cRNA was hybridized overnight to Human HT-12 V4 BeadChip arrays (IIlumina), which contains >47,000 probes, and scanned on an Illumina BeadStation to generate signal intensity values.

### TFA data generation

For the SLE cohort, a quantitative reverse transcription PCR platform was used. Globin reduced RNA was reverse-transcribed to cDNA using the High-Capacity cDNA Reverse Transcription Kit (Thermo Fisher Scientific), followed by specific target preamplification for 14 cycles in the presence of a pool of 136 primer pairs, including 8 reference genes (Supplemental Methods) (DELTAgene Assays, Fluidigm). Preamplified cDNAs were treated with Exonuclease I (New England Biolabs) to remove unincorporated primers and the preamplified cDNAs and detection assays were loaded onto a 96.96 Dynamic Array IFC (Fluidigm). Real-time PCR was run using EvaGreen dye (Bio-Rad) for detection on a BioMark HD System (Fluidigm). Analysis was performed using the Real-Time Analysis Software package (Fluidigm) to determine cycle threshold (Ct) values, using linear (derivative) baseline correction and auto-detected, assay-specific threshold determination.

For the pregnancy cohort, a NanoString assay was used. 100ng of total RNA was hybridized overnight (18 h) to target genes contained in a custom gene expression nCounter Plex2 for GEx NanoString Assay (Supplemental Methods), following the manufacturer’s Gene Expression Assay protocol. Enrichment of hybridized reporter/capture complexes and RNA target was carried out using SamplePrep Station and signal detection was carried out in an nCounter Digital Analyzer set for high-resolution scanning. NanoString data analysis guidelines were followed to carry out normalization to assay positive controls and to subtract background noise. Normalization to housekeeping genes included in custom gene panel (Supplemental Methods) was carried out using housekeeping-gene global geometric mean approach. Resulting normalized values were reported for downstream statistical analysis.

### Statistical Analyses

Two-group comparisons (t-tests) were run on log2 FC values between SLE and healthy controls to determine modules that showed significant differences between the two groups. For the longitudinal pregnancy data, mixed effects models, using the lme4 package in R [41], were run to compare pregnancy versus healthy controls over time. Principal component analysis (PCA) was performed on all healthy controls but no significant differences were found according to gender, age, or time of sample collection, so all healthy controls samples were included.

## Abbreviations

BIIR: Baylor Institute for Immunology Research
COPD: Chronic Obstructive Pulmonary Disease
CRP: C-Reactive Protein
Ct: Cycle threshold
FC: Fold Change
FDR: False Discovery Rate
GO_BP: Gene Ontology Biologic Process
GO_MF: Gene Ontology Molecular Function
HIV: Human Immunodeficiency Virus
IFN: Interferon
IRB: Institutional Review Board
KEGG: Kyoto Encyclopedia of Genes and Genomes
MS: Multiple Sclerosis
PCA: Principle Component Analysis
PCR: Polymerase Chain Reaction
PID: Primary Immune Deficiency
RF: Random Forest
ROC: Receiver Operating Characteristic
RSV: Respiratory Syncytial Virus
SLE: Systemic Lupus Erythematosus
SLEDAI: SLE Disease Activity Index
SOJIA: Systemic onset Juvenile Idiopathic Arthritis
TB: Tuberculosis
TBD: To Be Determined
TFA: Transcriptome Fingerprinting Assay

## Declarations

Ethics approval and consent to participate

Each of the studies contributing samples for this manuscript was independently approved by the BIIR IRB (IRB #’s 009-240, 006-177, 002-197, 009-257, H-18029, HE-470506, 011-173.

## Availability of data and material

Raw gene expression data was deposited in the Gene Expression Omnibus, https://www.ncbi.nlm.nih.gov/geo/, under the accession GSE100150.

## Authors’ contributions

Microarray data were generated at the Baylor Institute for Immunology Research and Benaroya Research Institute. qPCR data were generated at the Benaroya Research Institute. N.B. developed the modular framework. M.C.A, N.B, P.K, V.H.G, and D.C developed the TFA panel. M.C.A, P.K, and E.W analyzed data. L.C, N.J, J.T.P, and R.A.L contributed the samples run on TFA. J.T.P, G.K, M.F.L, H.B.R, A.O, M.B, C.B, M.L, R.W, C.G, G.L, M.E.C, J.S, F.K, K.P, A.M, O.R, V.P, and D.C contributed the microarray data for module generation. P.S.L, R.A.L, J.B, C.Q, S.P, and D.C.A provided support and contributed to study design. M.C.A and D.C wrote the manuscript.

## Acknowledgements

We thank Dr. G Nepom for helpful comments on the experimental design. We thank Quynh-Anh Nguyen, Kimberly O’Brien, Dimitry Popov, Michael Mason, and Cate Speake for technical assistance. We thank Brad Zeitner for development of software.

---

**Supplemental Figure 1:**
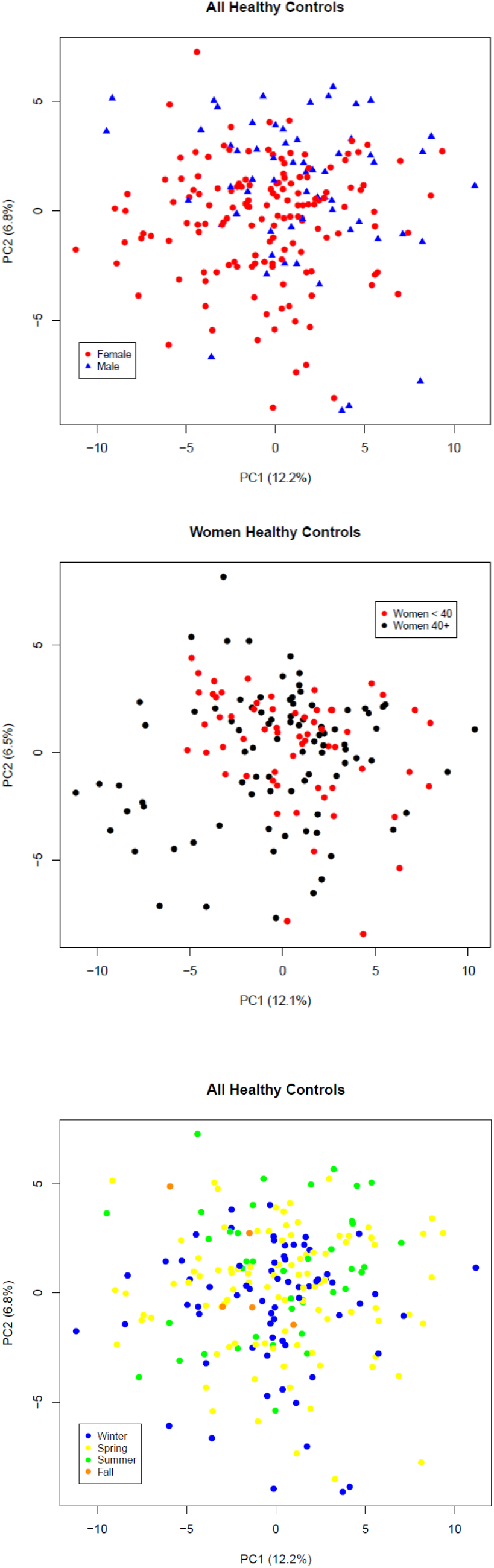
TFA gene expression in healthy controls. **(A)** Gene expression values from all healthy controls were used for principal component analysis. Scores from principal component 1 (PC1) and 2 (PC2) were plotted with these 2 components explaining about 19% of the variability in the data. There were no group differences attributable to gender among the expression values (red=female, blue=male). **(B)** Plot of PC1 and PC2 from principal component analysis of gene expression values for all female controls, which again explains approximately 19% of the variability in the data. There were no group differences attributable to age (as a surrogate for child bearing status) in the expression values (red=women <40yo, black=women>40yo). **(C)** No significant differences were found between healthy control samples obtained in different seasons as shown in this principal component plot (blue=winter, yellow=spring, green=summer, and orange=fall).

**Supplemental Figure 2:**
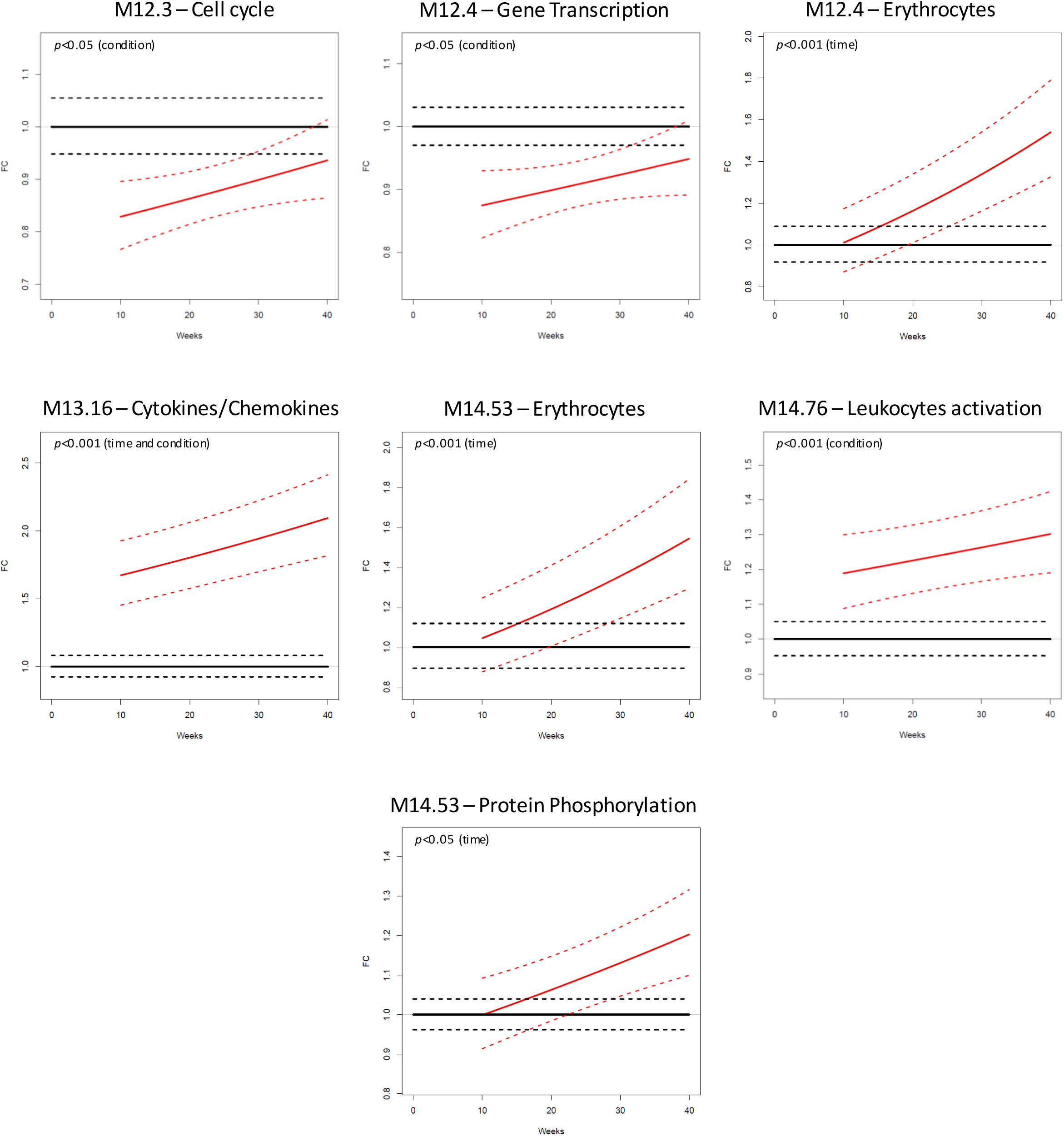
TFA assessment during healthy pregnancy demonstrates longitudinal immunological changes. Average FC expression levels referenced to healthy controls from the blood of 12 healthy pregnant women and 18 healthy non-pregnant controls. Shown are 7 of the 11 significant modules (see Figure 4 in the main text for the other 4). A linear mixed effects model was fit to the longitudinal data from pregnant women and healthy controls to determine if there was a trend over time for the pregnant women (shown as the time p-value) and if there was a significant difference between pregnant women and healthy controls over time (shown as the condition p-value).

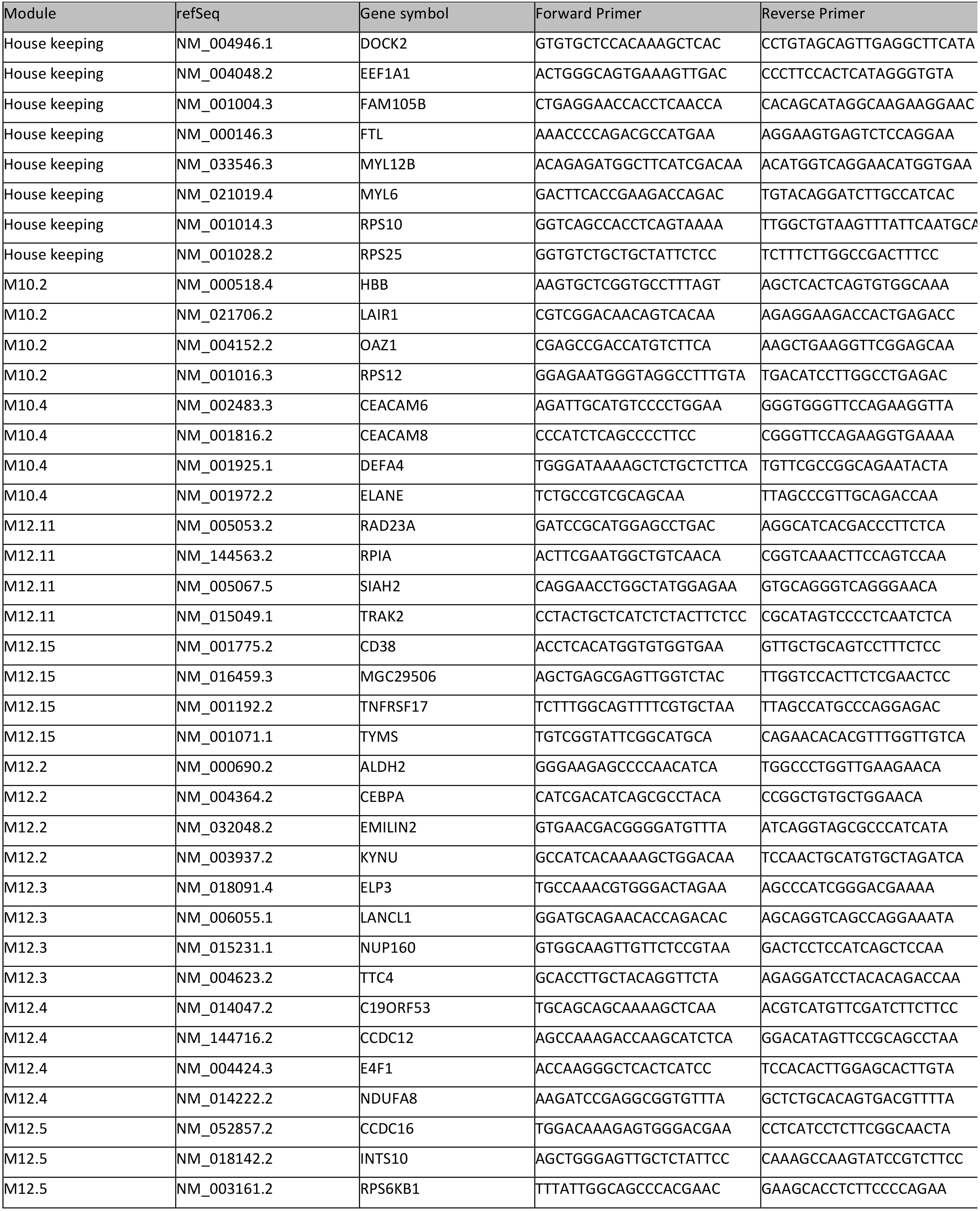
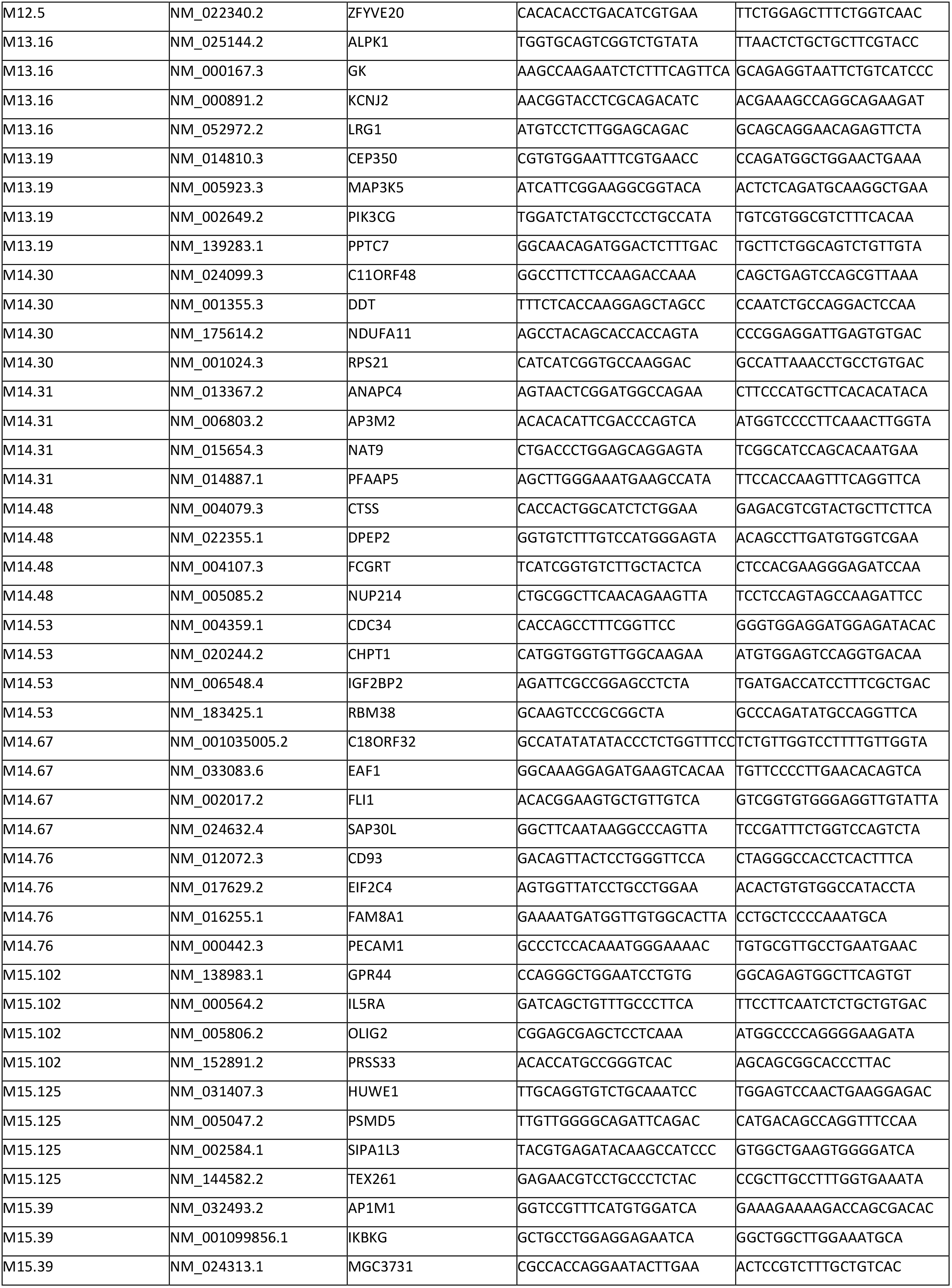
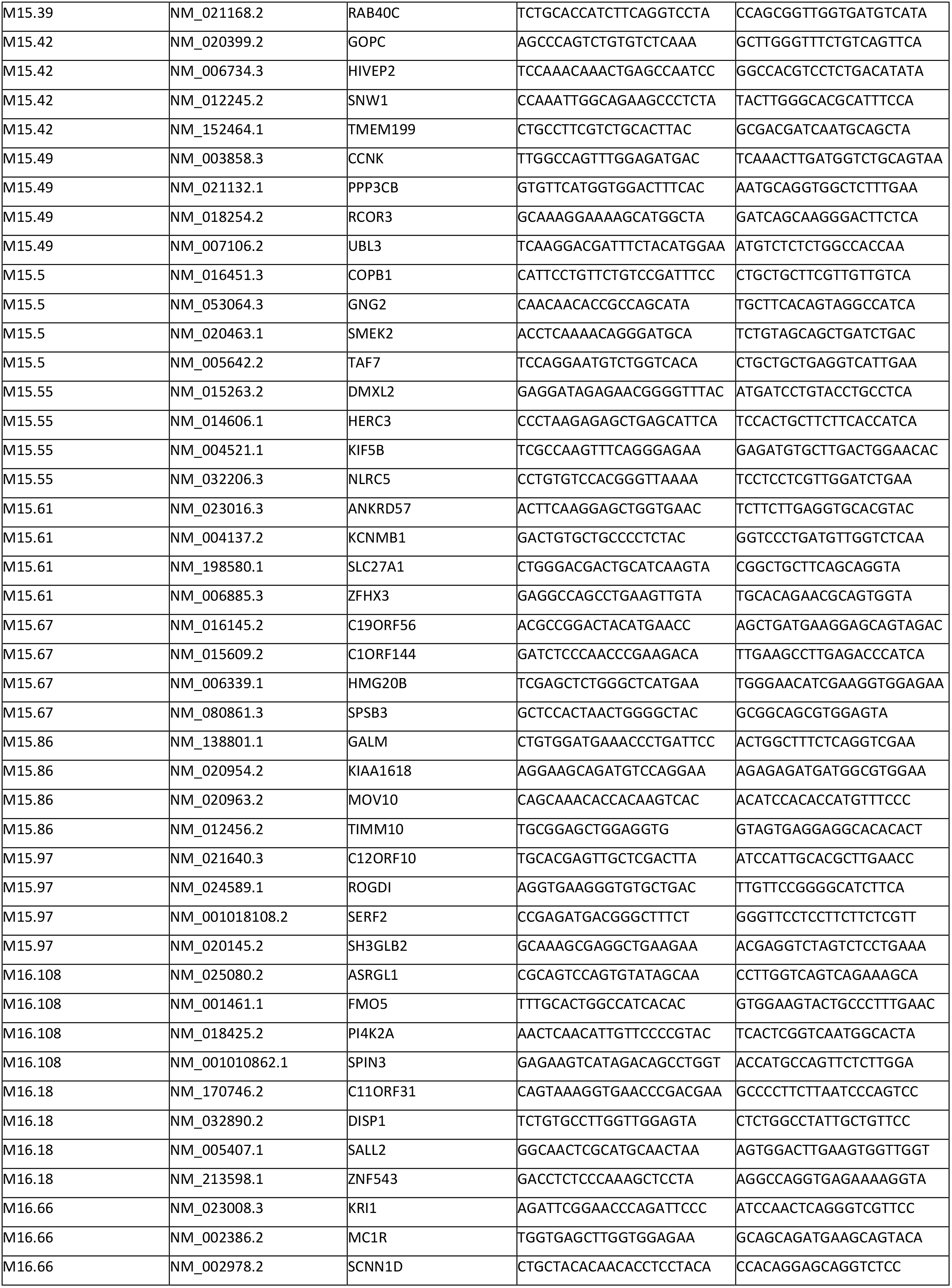
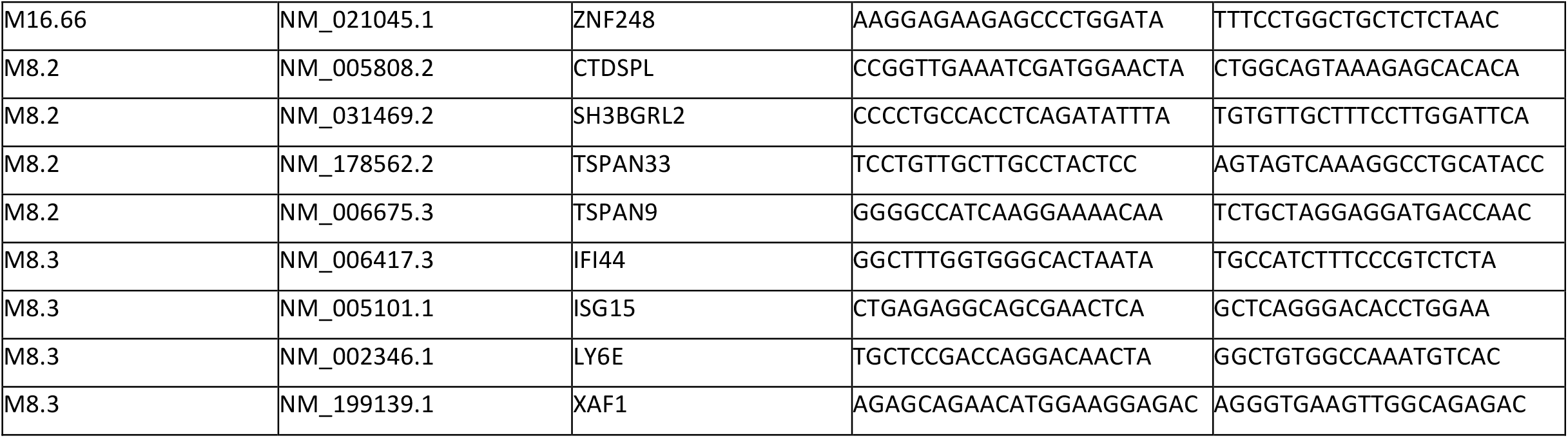
TFA primers. Listed are the primers used for the 8 housekeeping and 132 TFA genes.

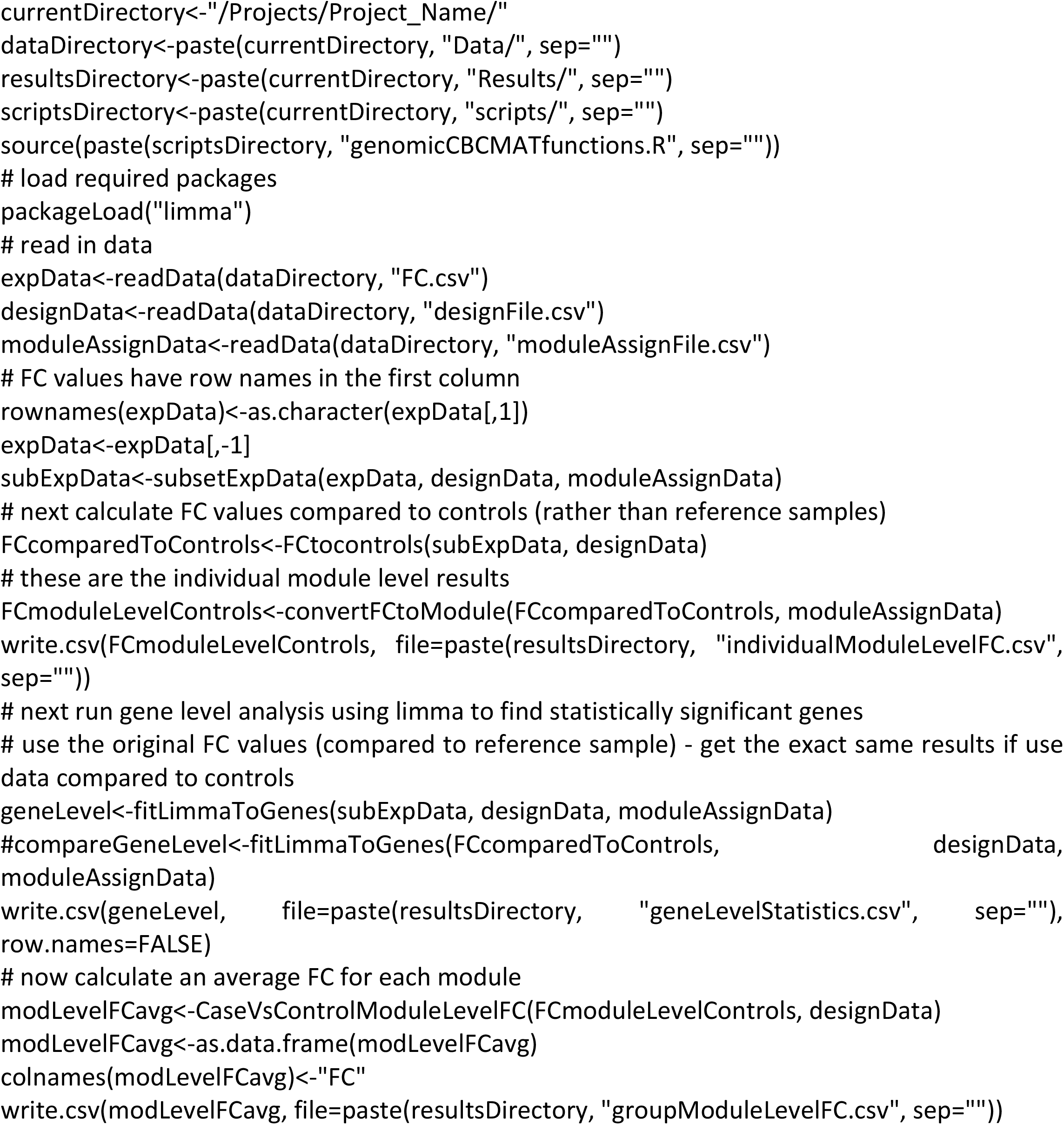
TFA Analysis Code. The R code below can be used to analyze TFA data that have been converted from Ct values to FC relative to reference samples. The structure expected for this analysis is a directory for the # set up the project directory

The functions that are called above are defined below:

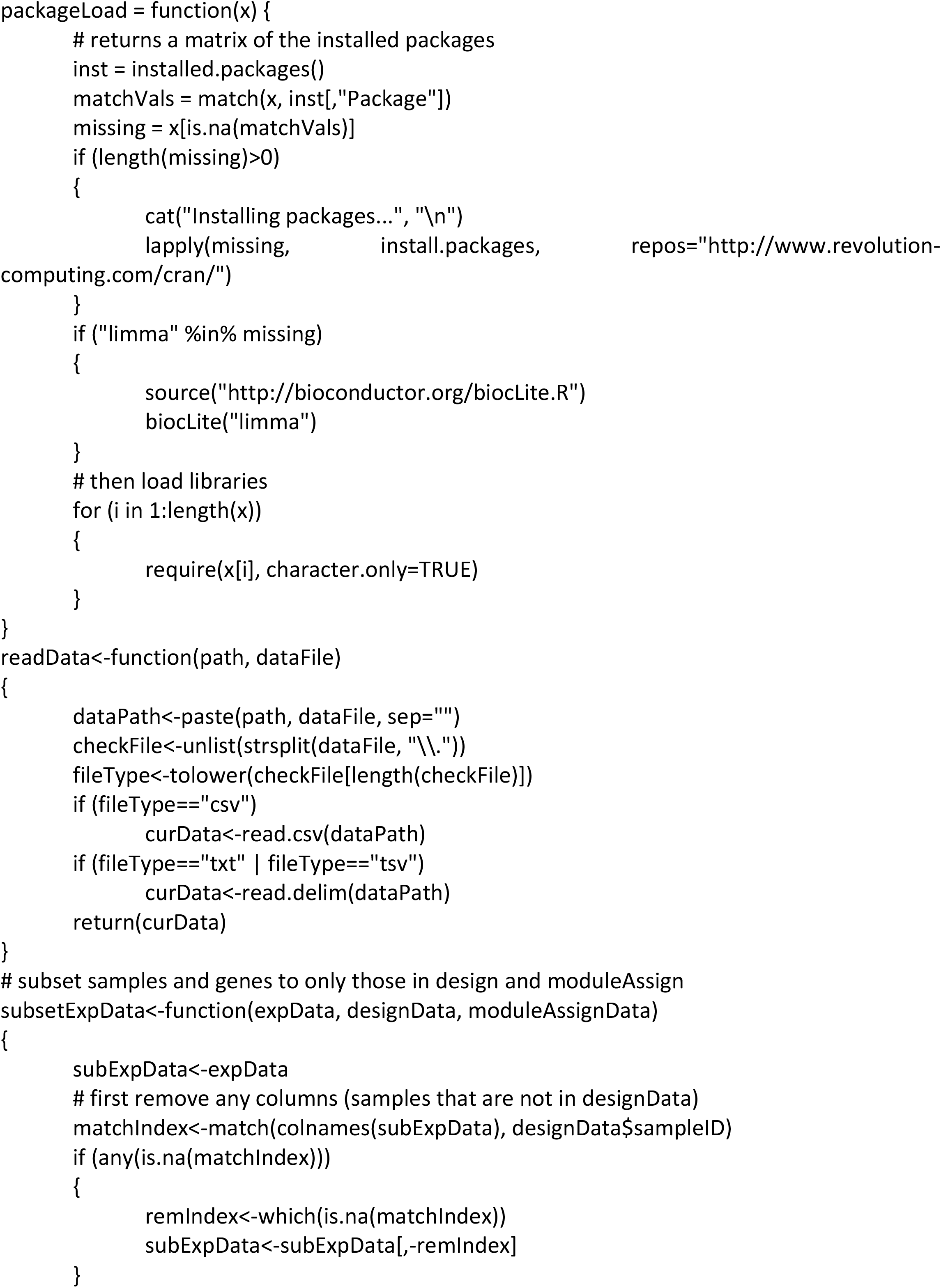
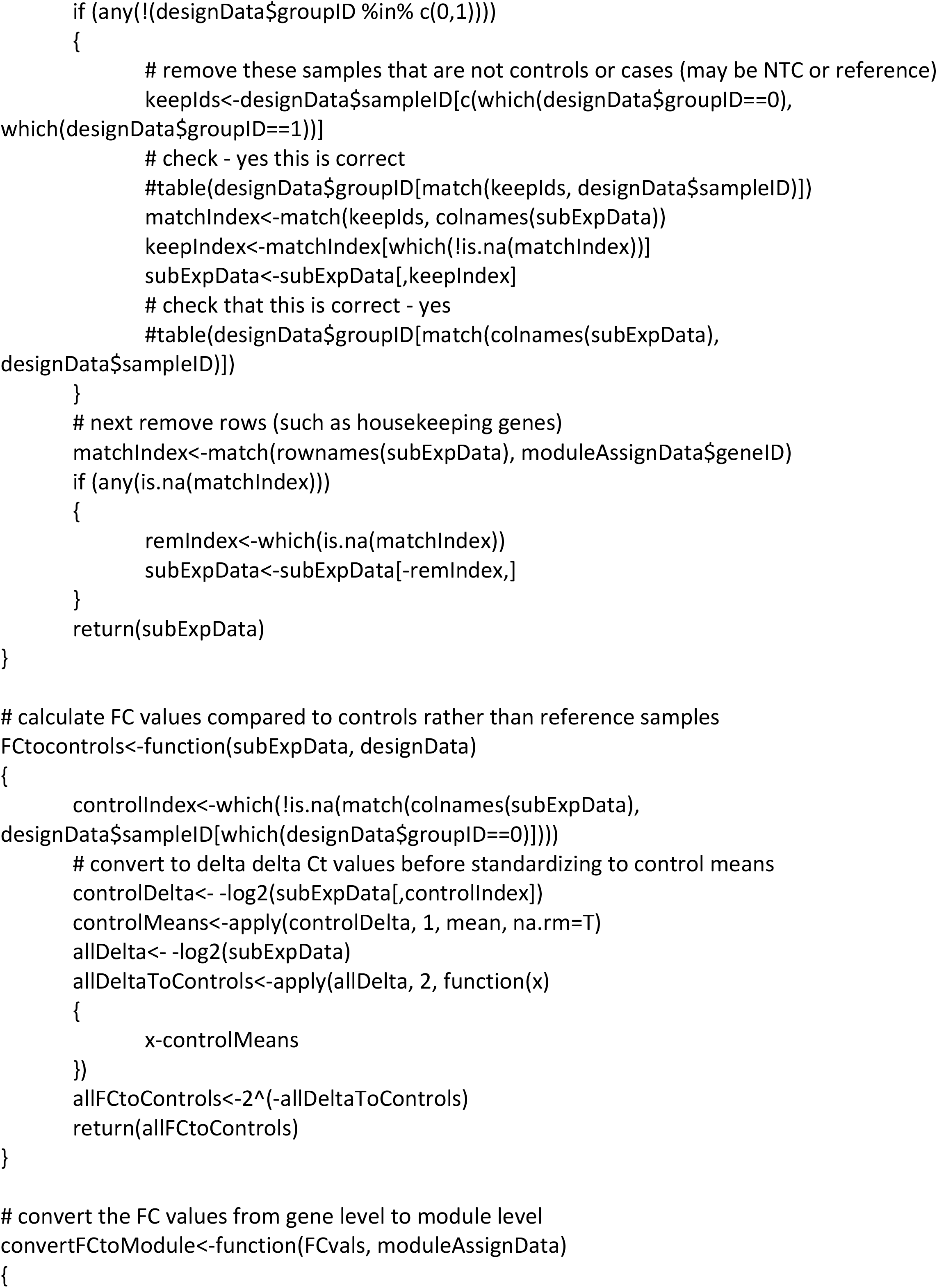
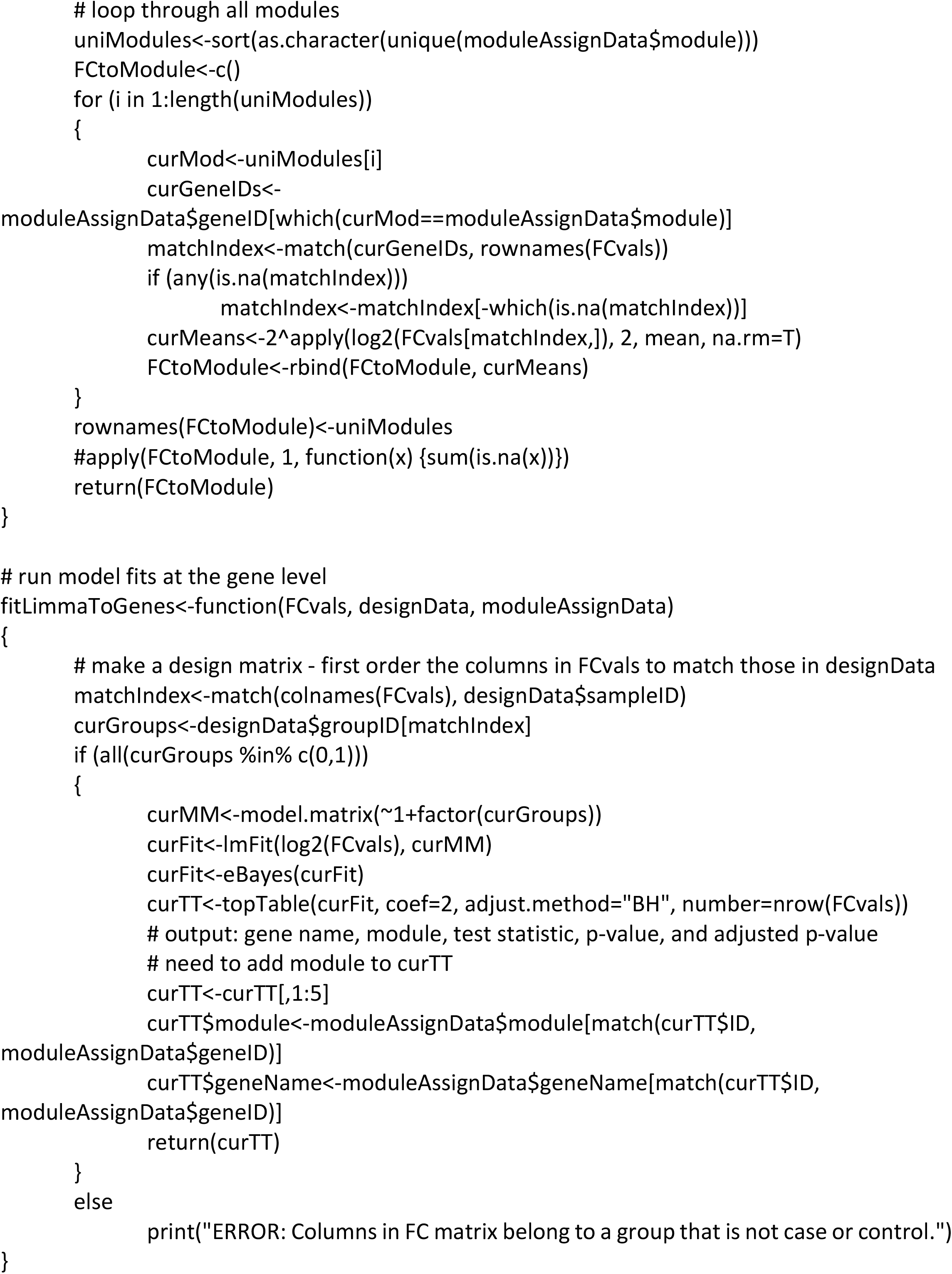
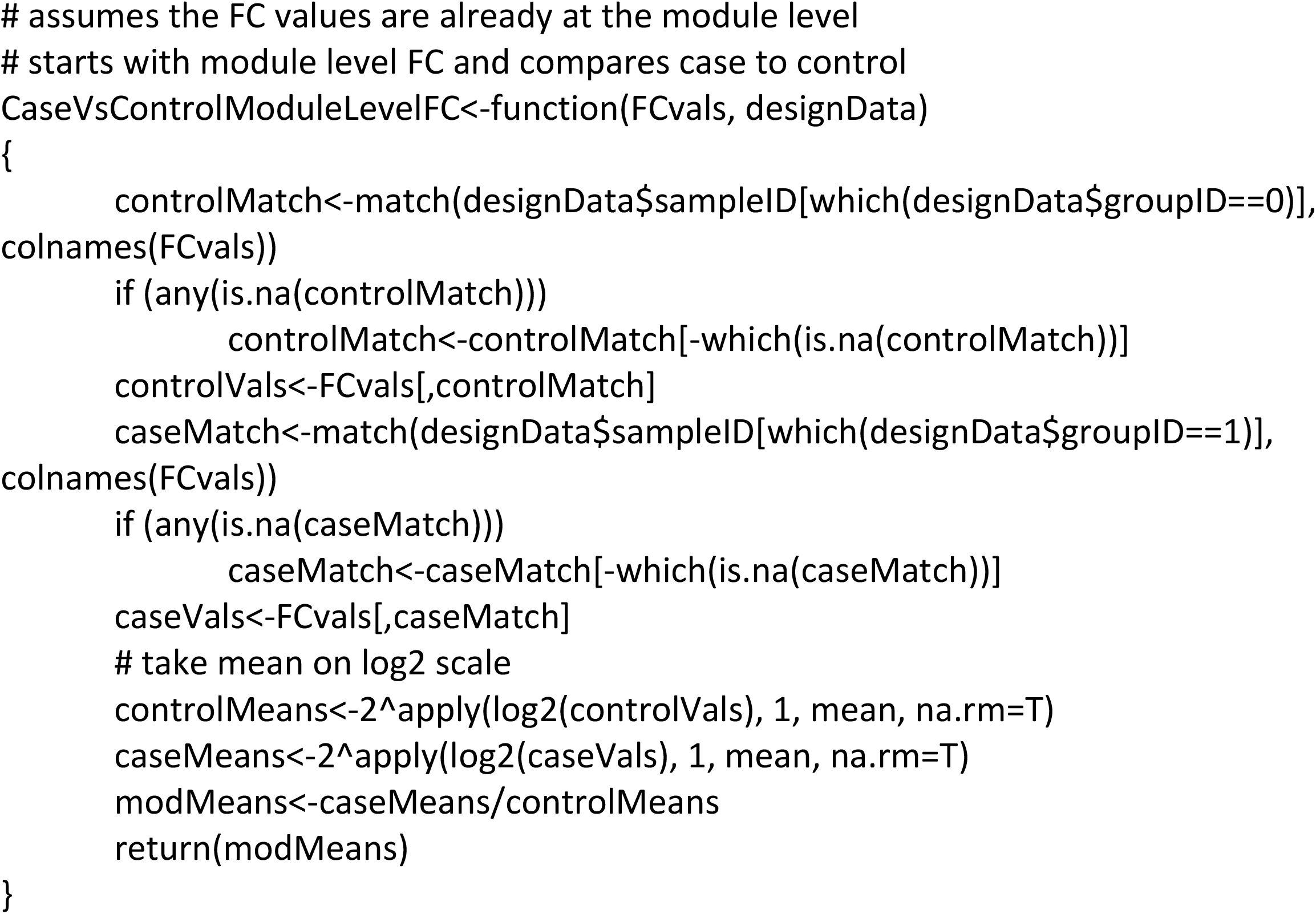

